# Chimpanzee play sequences are structured hierarchically as games

**DOI:** 10.1101/2022.06.14.496075

**Authors:** Alexander Mielke, Susana Carvalho

## Abstract

Social play is ubiquitous in the development of many animal species and involves players adapting actions flexibly to their own previous actions and partner responses. Play differs from other behavioural contexts for which fine-scale analyses of action sequences are available, such as tool use and communication, in that its form is not defined by its functions, making it potentially more unpredictable. In humans, play is often organised in games, where players know context-appropriate actions but string them together unpredictably. Here, we use the sequential nature of play elements to explore whether play elements in chimpanzees are structured hierarchically and follow predictable game-like patterns. Based on 5711 play elements from 143 bouts, we extracted individual-level play sequences of 11 Western chimpanzees (*Pan troglodytes verus*) of different ages from the Bossou community. We detected transition probabilities between play elements that exceeded expected levels and show that play elements form hierarchically clustered and interchangeable groups, indicative of at least six ‘games’ that can be identified from transition networks, some with different roles for different players. We also show that increased information about preceding play elements improved predictability of subsequent elements, further indicating that play elements are not strung together randomly but that flexible action rules underlie their usage. Thus, chimpanzee play is hierarchically structured in short ‘games’ which limit acceptable play elements and allow players to predict and adapt to partners’ actions. This ‘grammar of action’ approach to social interactions can be valuable in understanding cognitive and communicative abilities within and across species.

## Introduction

Animal lives take place in time – actions happen sequentially in response to changing environmental stimuli and the behaviour of other individuals. Particularly in social interactions, each action is a decision based on the social environment, the actor’s previous behaviour, the partner’s reactions, and intended outcomes (Bshary & Oliveira, 2015). Sequential social decisions are therefore an important window into the complexity of animal decision-making abilities (Gygax et al., 2021). Sequences can be considered complex for participants and bystanders if contingencies between actions are increasingly removed in time or with increasing flexibility and decreased determinism of transitions between actions (Kershenbaum et al., 2016). In many animal species, social play involves rapid exchanges of actions between several participants that often appear random to observers, making it one of the most complex social contexts individuals are involved in daily. Multiple individuals combine distinguishable and discrete actions (‘play elements’) in temporal patterns (‘play sequences’) and adapt to partners’ actions. This complexity can provide a unique window to unravel fast-paced decision-making in sequential exchanges between players. However, we currently lack a framework to understand how predictable or flexible play really is.

Sequential decision-making processes have been investigated in some detail in tool use and communication. In both, there is increasing evidence for predictable decision-making and hierarchical sequential structures. For example, New Caledonian crows (Hunt & Gray, 2004; Wimpenny et al., 2009) as well as several primate species (Boesch et al., 2020; Carvalho et al., 2008; Deblauwe et al., 2006; Estienne et al., 2017; Hihara et al., 2003; Martin-Ordas et al., 2012) use sequences of steps, often involving multiple objects, to solve problems using tools and tool sets. In chimpanzees, stone tool use (Carvalho et al., 2008; Sirianni et al., 2015), termite fishing (Deblauwe et al., 2006), and digging for underground bee nests (Estienne et al., 2017) have been analysed as complex sequences of individual decisions. Similarly, vocal patterns of bats (Bohn et al., 2009), birds (Berwick et al., 2011; Engesser et al., 2016; Sasahara et al., 2012; ten Cate, 2014), cetaceans (Allen et al., 2019), rock hyraxes (Kershenbaum et al., 2012), and primates (Arcadi, 1996; Arnold & Zuberbühler, 2008; Clarke et al., 2006; Girard-Buttoz et al., 2022; Leroux et al., 2021; Ouattara et al., 2009) have been described as temporal sequences with different degrees of predictability, combinatorial complexity, and hierarchical structure. This has often been related to the evolution of syntax (Zuberbühler, 2019). Increasingly, communication sequences are found for other communicative modalities, such as gestures and facial signals (Aychet et al., 2021; Genty & Byrne, 2010; Graham et al., 2020; Liebal et al., 2004; McCarthy et al., 2013; Safryghin et al., 2021). Studies have shown that Markov processes (i.e., elements are predicted by a finite number of antecedent elements) are insufficient for describing vocal sequences (Kershenbaum et al., 2014). Different species show turn-taking in exchanges and adapt their signals as sequential response to a partner’s actions (Demartsev et al., 2018; Fröhlich, 2017).

In both tool use and communication research, the form of sequences is defined partially by their function: in tool use, an ‘optimal’ sequence exists that allows individuals to access a resource (Estienne et al., 2017). In communication, complexity is limited by the need to be understood, which cause sequences to be predictable and short. Songs are not constrained the same way, often containing hundreds of hierarchically structured elements (Berwick et al., 2011). Given that most species do not create song-like vocalisations, understanding sequences in social interactions (their ‘grammar of action’; Pastra & Aloimonos, 2012) could potentially allow for a broader perspective on action sequences. Play is a prime candidate because the form of play is not necessarily the results of a specific function – play has been hypothesised to have evolved as practice for future challenges facing individuals, so it is defined by its unpredictability compared to ‘real’ interactions (Fagen, 1981; Palagi et al., 2004; Smith, 1982).

Play behaviour, at least during some parts of development, is common in most mammals and birds (Diamond & Bond, 2003; Fagen, 1981), and exists in some reptile, fish, and amphibian species (Burghardt, 2015) and in octopuses (Kuba et al., 2006), indicating that it is an ancient behavioural context. Species can have large repertoires of distinct elements (Petrů et al., 2009). Play signals are deliberately used to prevent play from breaking down when intentions are unclear or risk is high (Cordoni & Palagi, 2012), and extend the length of play bouts (Waller & Cherry, 2012). We have yet to learn how coordinated other play actions are, and whether expected responses to certain action sequences are socially learned or innate. In human play, there are specific, socially learned arbitrary rule systems that govern what we call ‘games’ (Leisterer-Peoples et al., 2021): in a game, certain actions and sequences are allowed or not, but their order can be flexible. For example, in hide-and-seek, hiding is allowed, but laughing loudly is counterproductive, where and how to hide is up to the player. There is evidence that apes have standardised games and play them with each other and human partners (Costa et al., 2019; Pika & Zuberbühler, 2008; Tanner & Byrne, 2010) – however, these examples focus on special contexts (e.g., playing in water, playing socially with objects), and we do not have a method to determine how widespread predictable behavioural rules are.

The Bossou Western chimpanzees (*Pan troglodytes verus*) have been studied since 1976 (Matsuzawa & Humle, 2011). An ‘outdoor laboratory’ was created in 1988 as a clearing in the territory of the community where stones and nuts are provided to study tool use, with standardised video recordings available for over 30 years. Because the chimpanzees spend considerable time there, social and object play can be observed regularly (Myowa-Yamakoshi & Yamakoshi, 2011). In chimpanzees, infants and juveniles play more than older subadults and adult individuals (Cordoni & Palagi, 2011), but chimpanzees are among the few species where adult play seems common (Fernandez-Duque et al., 2000) and fulfils several functions, especially in conflict regulation and stress reduction (Palagi et al., 2004). Chimpanzees play with and without objects (Koops et al., 2015), and solitary and socially, often involving more than two players (Cordoni et al., 2018; Shimada, 2013). Play signals are used to advertise willingness initiate play bouts and increase their duration (Davila Ross et al., 2009; Matsusaka, 2004; Waller & Dunbar, 2005), and there is good evidence that chimpanzees show matching or mimicry of partners’ play face and laughter (Davila-Ross et al., 2011; Ross et al., 2014). Gestures can occur in sequences during play (Bard et al., 2014), especially if partners fail to respond initially, with tactile and audible gestures usually occurring early in the sequence (McCarthy et al., 2013) and younger individuals producing more tactile gestures (Fröhlich et al., 2016). The cooperative and coordinated nature of play (multiple individuals adapting their behaviour in real-time to sustain the interaction) has been used to study higher socio-cognitive skills such as joint intention and shared intentionality with varying results (Bekoff & Allen, 1998; Pika & Zuberbühler, 2008; Tomasello et al., 2005), and joint commitment and joint action (Heesen, Bangerter, et al., 2021; Heesen et al., 2017; Heesen, Zuberbühler, et al., 2021). Anecdotal evidence from the Bossou chimpanzees has repeatedly indicated that chimpanzee play might involve aspects of pretence or imagination (Matsuzawa, 2020; Nakamura, 2012). Our focus is on the form of play, how elements are strung together, which has its own implication for cognitive evolution.

For this study, we tested whether sequences of play elements are predictable for players or contain a large amount of randomness, and whether we can identify hierarchical structure in sequence patterns. To do this, we ask two main questions: if I know the previous action (’antecedent’), can I predict the subsequent action (’consequent’)? And are there higher-order connections between elements, in the form of network clusters of interchangeable elements? This last aspect would indicate the presence of ‘games’: once we are playing a game, certain elements are permissible, but their order and exact usage can vary. This study specifically looks at transitions within individuals - partner behaviour is considered ‘noise’. This will reduce predictability, because actions that appear ‘unexpected’ here are possibly expected responses to partner actions. We hypothesize that some play elements are consistently more likely to follow specific antecedents than would be expected at random. Using the probabilities of each element and each transition to ‘predict’ which element will appear next, we expected classification accuracy that exceeds random assignment, and that higher-order sequences (AB, rather than B alone, to predict C) further improved prediction accuracy. We also hypothesized that, like communication in some species (Allen et al., 2019), we can detect hierarchical structures in transition networks (‘games’) as clusters of elements that are often used together and can be used interchangeably. Using the transition probabilities of each element to each other element, we can identify clusters of elements that have similar transition patterns (i.e., act like ‘synonyms’). The network structure allows us to identify elements that were essential to a game (in the sense that they occurred at higher rates than other elements in the cluster and connected other elements in the sequence; Carvalho et al., 2008). Lastly, we predict that the similarity and transition clusters overlap – i.e., we have clusters of elements are interchangeable and tightly linked in time.

## Methods

### Sample

We scanned 116h of video material from the Bossou video database (Matsuzawa & Humle, 2011), collected between 2009 and 2013. While footage from the Bossou outdoor lab has high video quality and filming consistency, the social composition of the group limits generalisability. The Bossou community at the time was small (around 13 individuals) (Matsuzawa & Humle, 2011). Due to the age distribution, there was only one infant, one juvenile, and one subadult individual in the group during data collection – making it difficult to differentiate between age effects and individual preferences (Fröhlich et al., 2016). Eleven individuals were observed playing at least once; however, the distribution of observations was highly skewed, with the two juvenile/subadult players each participating in about 75% of all play bouts, while none of the adults participated in more than 20% of play bouts. Thus, most play elements and transitions were provided by two individuals, often playing with each other. In this study, we do not control for individual or age differences in play behaviour and sequences, due to the limited sample. These could make play transitions more predictable (individuals or specific age groups might have standardised ways of reacting that other group members know). Considerably more data would be necessary to control for individual- or dyad-level effects in transition patterns. We identified 143 bouts of social play across 35 videos - defined as play involving at least 2 individuals, with a new bout started if both individuals stopped playing for at least 5 seconds continuously. Bouts consisted of between 3 and 181 individual play elements (mean = 30.3), including between 2 and 4 players at any given time. For analyses, the bouts were split into individual-level bouts (every play element an individual performed during a bout), resulting in 306 individual-bouts.

### Coding Scheme

The coding scheme, with detailed definitions of all play elements and coding conventions can be found in the associated repository. Potential play elements were identified from several sources – primarily, every behaviour indicated in Nishida et al., (2010) as potential play behaviour, the literature on ape gestural repertoires (Genty et al., 2009; Graham et al., 2017; Hobaiter & Byrne, 2011, 2014), previous chimpanzee play literature (Fröhlich et al., 2016), and descriptions of play elements in primates more widely (Petrů et al., 2009). Often, these sources use different terms for similar play elements, and the definitions used here do not always overlap perfectly with those used previously. To our knowledge, the ethogram used here is the most detailed ethogram for chimpanzee play to date. Play elements can roughly be categorised as contact or non-contact, and as events (countable, one-off or repeated actions) or states (continuous behaviour with a clear start and end point). Social object play formed its own category, with multiple different ways of interacting with detached objects (mainly stones, nuts, and sticks) available. In total, our ethogram contained 118 different play elements, of which 106 were observed at least once. We assumed that the elements we defined are meaningfully different from each other. This might not be the case: the difference between *Retreat* (walking away from partner), *Flee* (running away from partner), and *Retreat Backwards* (walking away from partner while looking at them) might be an artifact of the coding scheme.

Coding was done using BORIS v.7.9 video coding software (Friard & Gamba, 2016). We coded bouts one player at a time and marked the start of every change in play element and mark all active play elements at that time point. For example, if an individual goes *bipedal*, this is marked. If, while bipedal, the individual approaches the partner, we would mark *bipedal/approach*. If they would then raise their arm while performing those actions, we would mark *bipedal/approach/arm raise*, and so on. This leaves us with a string of play elements with a time stamp for initiation. If any player stopped playing (i.e., no play element was active), a Break was coded. The duration of play elements was available but was not considered in this study – we focus entirely on the sequential order.

Video coding of entire play bouts is slow, due to fast changes of behaviour and movements, and researchers usually focus only on play initiation and re-initiations (Heesen, Bangerter, et al., 2021; Hobaiter & Byrne, 2011). Due to the challenges of this detailed coding approach, no inter-rater reliability was performed, and results must be viewed with this limitation. Predictability should be higher in studies using simpler coding schemes, so if we can show high predictability using the current ethogram, we have taken the conservative approach. The dataset currently contains 5711 play elements. Where possible, we present results including uncertainties, and used permutation and bootstrapping approaches to discriminate between spurious and reliable transition patterns.

### Pre-processing

All pre-processing and analyses were conducted in R statistical computing software (R Development Core Team & R Core Team, 2020). The video coding data needed pre-processing to deal with three main problems inherent to the coding process: rare elements; some artificially common elements; and establishing the sequential order of co-occurring elements.

To robustly establish probabilities of transitions between elements, rare elements are a problem (Silge & Robinson, 2017). For example, if an element only occurs three times, and each time transitions into a different element, we do not know if the high transition probability would disappear with increasing sample size. We set the threshold at 20 occurrences per play element. However, removing these cases completely (as is often done in linguistic studies; Silge & Robinson, 2017) would be wasteful given the sample size of this study. For most play elements, we defined *a priori* with which other play element they would be combined if too few occurrences were observed (see associated repository). Replacement elements were chosen based on similarity of movement. If the combination after this lumping process failed to reach the threshold, we nevertheless retained it. Thus, our rarest element had 9 occurrences (see associated repository for occurrence probabilities of all play elements before and after pre-processing). After this step, 68 play elements remained.

Some elements occurred at much higher frequencies than others. The seven most common elements (Bipedal, Hold, Follow-Other, Approach, Retreat-backwards, Retreat, Flee) were all coded continuously and therefore were noted every time a change occurred while they were active. Imagine a musical piece on the piano: sometimes one note is held while others are played. In play, a chimpanzee could go bipedal, but then perform other actions while the *Bipedal* was marked at every change in event. These elements potentially skew transition probabilities and mask transitions between other elements. Ideally, we want a sequence that reflects when individuals made the choice to use a specific element. We addressed this by detecting cases where one of those seven elements occurred multiple times in a row, and only retained the first case. If players stopped the continuous action (e.g., stopped fleeing, then started again), the element was counted again.

In play, it is possible to go *Bipedal*, *Arm Swing* with one arm and *Hit* the partner with the other arm. This is problematic in terms of the transitions - does *Bipedal* lead to *Arm Swing*; or *Arm Swing* to *Bipedal*? This problem also occurs mainly because some elements (e.g., *Flee* or *Bipedal*) are continuous states, while others (e.g., *Kick*) have a clearly defined beginning and end. We used permutations – randomly assigning order within co-occurring elements and repeating all analyses 1000 times with different orders – as there was no *a priori* reason to assign primacy to one co-occurring element over another. Thus, all described transition probabilities are averages over multiple permutations, which is why transition counts are not integers. Two alternative approaches (random sampling of only one of the co-occurring elements, bag-of-words) can be performed using the attached R scripts and generally showed similar results.

### Transition Probabilities

The transition probability between antecedent and consequent were defined by the number of times the consequent followed the antecedent, divided by the number of times any element followed the antecedent (conditional probability). The antecedent could be a single element (used to establish first-order n-grams, networks, and transition similarities), but also n-grams of different order (e.g., first order: *Hit*; second order: *Hit*/*Slap;* third order: *Hit/Slap/Tickle* etc). The latter approach was taken to determine whether increased information about antecedents increases prediction accuracy. Current sample size prevents us from analysing long sequences, as the number of possible transitions increases exponentially with each new level. We limited the analyses to a maximum of 3 antecedent elements. We restricted ourselves to one-element consequents and did not consider non-adjacent contingencies (Sonnweber et al., 2015).

The large number of possible combinations combined with a small dataset and the small number of individuals leads necessarily to overfitting: some combinations will only occur a few times and adding new information could influence our understanding of their function. We did two things to counter this: rare elements were combined, as described above. Where possible, we report some measure of robustness to give the reader an understanding of how reliable results were. Robustness was established using bootstrapping procedures – randomly selecting 1,000 subsets of the data and establishing transition probabilities within those subsets.

### Randomisation Procedures

To test which elements followed which antecedent, we created a null model of ‘expected’ transitions using permutations of observed patterns. We chose this resampling approach over collocation analysis (Bosshard et al., 2021) to account for the regular co-occurrence of play elements that is usually not seen in single-modularity communication. We repeatedly randomized the order of elements across bouts: while the number of elements per bout, the probability of elements to occur across bouts, and the position of Breaks and missing data in each bout were kept the same, we randomly assigned element positions. Thus, transitions are considered significant if they were observed more often than would be expected if play elements were just strung together randomly given their base probabilities. We ran 1000 randomisations to create the expected distribution for each transition and compare whether the observed transition probability fell within this distribution or not. To compare the observed and expected values, we provide a p-value (how many of the 1000 randomisations show higher transition probabilities than observed; Mielke et al., 2021). We report transitions that occurred at least five times and that were significant at 0.01 level (i.e., the observed value was higher than for 990 permutations). These calculations also constituted the basis for the network clusters described below.

### Prediction Accuracy

To understand the predictability of transitions rules, we applied the probabilities derived from a subset of the data to ‘unknown’ test data and explored how well the former predicted the latter (Chollet & Allaire, 2018). We tested the predictability of elements within bouts by calculating transition probabilities for 95% of all other bouts, then predicting each element in the remaining 5% of bouts based on their antecedents (k-fold validation). This was repeated 1,000 times per bout. We tested the expected correct classification if the consequent element was only determined by base occurrence probabilities (null model). The difference between this value and the observed prediction accuracy of the models tells us how much knowledge of the antecedent increases our predictions. Aside from using one element as antecedent (describing a simple Markov process), we repeated the process with two or three elements as antecedents (n-gram prediction; (Eisenstein, 2019). For higher-order antecedents, the probabilities of the lower-order antecedents were combined (interpolation) – therefore, for *Approach*/*Stare At*/*Hit* as third-order antecedent, the probability is the product of the probabilities of the triad, *Stare At*/*Hit*, and *Hit*. This was done because many higher-order antecedents only occurred infrequently, and no information would otherwise be available as to which consequent was appropriate. For transitions that were never observed, Laplace smoothing was applied, assigning them one occurrence, and adapting all transitions accordingly (Eisenstein, 2019). If the prediction accuracy under those conditions was higher than for one element, this indicated hierarchical processes - for example, if *Hit* correctly predicts to *Hold* 10% of the time, but *Stare At*/*Hit* leads to *Hold* in 80% of the time, then the sequence order added information. We present the mean correct classification rate across all bouts and elements. In addition to predictions based on the transition probabilities, we implemented a naïve Bayes classifier using the ‘e1071’ package in R (Meyer et al., 2021). Naïve Bayes classifiers use vectors of feature values (in our case, the previous play element, two previous play elements, etc.) to predict the correct consequent using Bayes theorem (Eisenstein, 2019). Using an established classifier offers the advantage that classification is optimised and faster than the above-described prediction based on raw transition probabilities. However, naïve Bayes classifiers make a strong independence assumption, effectively assuming that the antecedents are independent from each other given the consequent class (Eisenstein, 2019). Therefore, while increased performance of the classifier with increasing number of antecedents would indicate that information about previous play actions increases predictability of what happens next, performance cannot be interpreted as based on sequential information.

### Similarity

We determined whether there were play elements that resembled each other in which elements followed them and tested whether we could find clusters of similar elements. This is similar to the identification of synonyms in language (Levshina, 2015), and we did it both to test whether our assignment of distinct elements during coding was meaningful and to see whether there were clusters of interchangeable elements. Each element was represented by a vector of transition probabilities with all elements. We applied Uniform Manifold Approximation and Projection (UMAP; McInnes et al., 2018) to achieve two-dimensional representation for each vector using the ‘umap’ package (Konopka, 2022). We established similarity between play elements by calculating the Euclidean distances between UMAP projections. To identify the optimal number of clusters for the hierarchical clustering, we used K-Means clustering as implemented in the ‘cluster’ R package (Maechler et al., 2022) to determine a) the optimal number of clusters, and b) the quality of the cluster solution. We present the silhouette value (Rousseeuw, 1987) to detect the best cluster solution, indicating an acceptable distance between clusters and coherence within clusters – any solution above 0.3 can be considered to show that there is more similarity within than between clusters. As cluster solutions differ based on the outcome of the UMAP dimension reduction, we repeated the dimension reduction and cluster detection 50 times with varying numbers of epochs for the UMAP (on average 7000 epochs) and continue using the most likely cluster solution. We plot the dendrogram for the optimal cluster solution and saved cluster memberships for later comparison with network clusters.

### Networks

Networks can be useful tools to visualise the connections between elements in communication networks and to identify clusters of elements that have above-expected connections with each other (Allen et al., 2019; Aychet et al., 2021; Barceló-Coblijn et al., 2017; Mielke et al., 2021; Weiss et al., 2014). Here, we created a network using all play elements as nodes and the transition probabilities between them as edges (Newman, 2010). Only transitions that were significantly more likely than expected and occurred at least 5 times, to make the network intelligible despite the large number of elements and ensure biological relevance. Edges were weighted, representing the transition probabilities between elements; and directed, meaning that each dyad of elements was represented with two values (A to B, B to A). We used the ‘igraph’ and ‘ggraph’ R packages (Csardi & Nepusz, 2006; Pedersen, 2021) to create and visualise networks. To test whether distinct ‘clusters’ of play elements existed in the network (indicating groups of play elements that have strong connections with each other but weak connections to the outside), we used the ‘cluster_optimal’ community detection algorithm in igraph, which maximises modularity of clusters (Csardi & Nepusz, 2006). Clusters were considered to represent stronger connections within than between clusters if the modularity value of the cluster solutions was larger than 0.3. Cluster solutions were compared to those produced by the similarity measure above.

## Results

### a) Non-random transitions

There were 1622 transitions that were observed at least one time. The histogram (Fig. 1) shows that most elements are followed by several different *consequents with low probabilities.* In only 4 cases did a consequent constituted more than 30% of all possible transitions of an *antecedent, with two of those (Drum Tree and Kick Dirt) being loops – the element was repeated sequentially*. At the same time, each element was observed to be followed by between 7 and 53 elements. Thus, there was no tight coupling between any two elements. This might indicate random assignment - any elements could be followed by any other. However, it might also mean situation-specific responses that were tailored to the players’ own previous action and the partners’ reaction, or predictability at a higher order (e.g., based on multiple antecedent).

**Figure 1:**
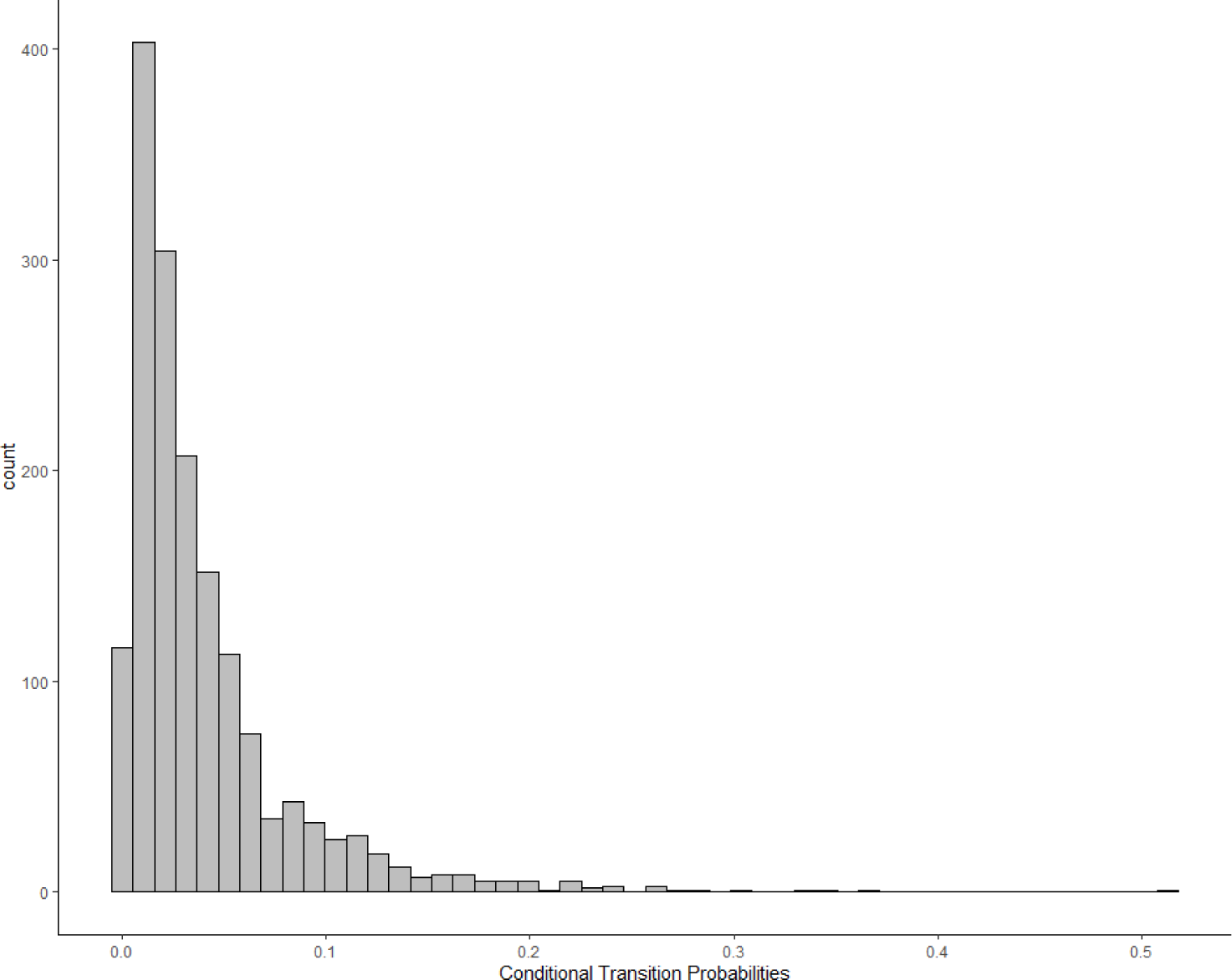
Transition probabilities for each simple antecedent - consequent pair

We also visualize how robust transitions were (Fig. 2). Using bootstraps, we created an interval around the observed transition probabilities. We plotted the range of values for each transition for the 1,000 bootstraps (calculated as the highest transition probability minus the lowest transition probability of A to B in the set) against the number of times the antecedent was observed. For some rare elements, transition probabilities remained volatile. Transition probabilities of rare elements therefore must be interpreted with caution, and elements will be filtered to exclude rare transitions – in all descriptions of ‘significant’ transitions and in the networks, only transitions that occurred at least 5 times were considered and reported.

**Figure 2:**
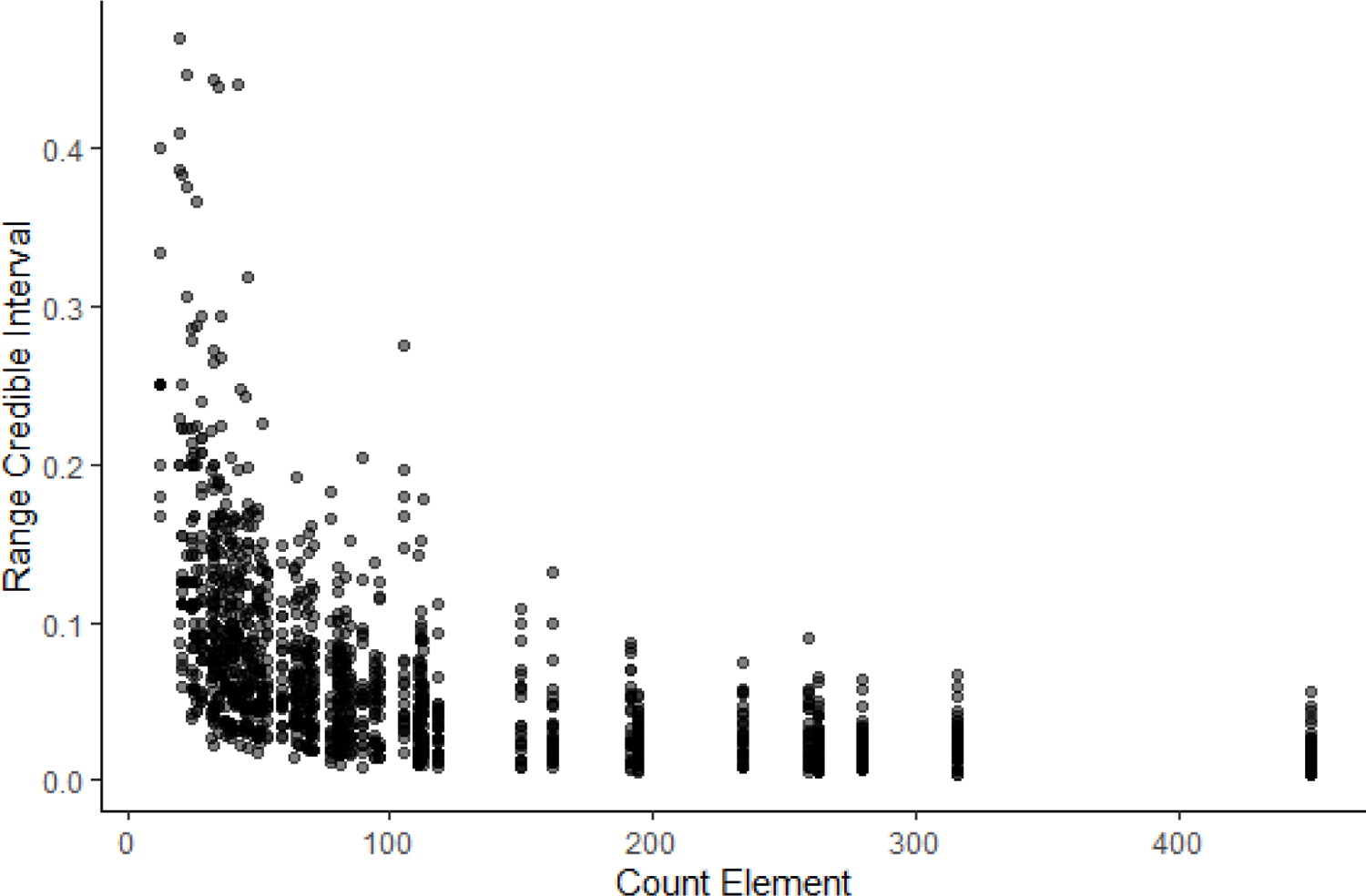
Range of bootstrapped transition probabilities compared to the occurrence of the antecedent. Transitions of rare antecedents are volatile.

In total, 146/1622 transitions (9 %) were significantly more likely than expected. More detailed depictions of these patterns can be seen in the network below and in the associated repository. When analysing the non-random transitions in detail, we found that many elements significantly followed themselves (21 out of 147 significant transitions). Several of the elements used here – for example, rocking or drumming on an object – are repeated actions and each occurrence was marked as independent event. In contrast to all observed transitions described above, many elements (17/68 elements) had no significant consequent, 14/68 had only one significant consequent, with the maximum number of significant transitions in one antecedent being 10 consequents (for *Holding* the partner and *Bipedal*).

### b) Next-element predictions

When applying the transition probabilities as predictions, increased information about antecedents increased predictability (Tab. 1). The basic probability of correctly predicting an element based on its occurrence probability (zero-order) was 0.03. By applying the probability of one antecedent (unigram; e.g., *Hit*) we increased the probability to 0.06 – almost a doubling of correct classification. When adding two antecedents (bigram; e.g., *Bipedal/Hit*), there was another rise to 0.11 – again, almost a doubling of correct classifications, and almost four times higher than having no information about antecedents. At the third order, we do not achieve further improvement. For the naïve Bayes classifier, using a more optimised approach that however assumes independence of antecedent elements, we achieve correct classification results of 0.09 as baseline, 0.13 for the first order, 0.25 for the second order, and 0.31 for the third order. Thus, additional information about preceding elements improved prediction accuracy. However, there was still a lot of unexplained variation.

**Table 1:** Correct prediction ability of consequent elements based on antecedents of different orders for the probability distribution and naïve Bayes classifier

| Order | Antecedent Example | Accuracy Probability | Accuracy Naïve Bayes |
| --- | --- | --- | --- |
| 0 | - | 0.030 | 0.086 |
| 1 | Hit | 0.057 | 0.153 |
| 2 | Stare At/Hit | 0.105 | 0.247 |
| 3 | Approach/Stare At/Hit | 0.105 | 0.308 |

### c) Similarity between elements

In Figure 3, we can see the dendrogram representation of hierarchical clusters of distances between transition probability vectors of all play elements. Elements connected through shorter branches and assigned the same cluster membership (same colour of branches) are considered more similar than those further away and with different colours. The best cluster solution, with silhouette value of 0.68 (indicating a well-distinguished cluster solution) contained 12 clusters. The cluster allocation can be seen in Table 2, and we will discuss their potential classification together with the network. What we can see here is that there were many elements that were similar in consequents. For example, *Kicking* the partner and *Jumping on* them were close, indicating that they could have been defined as a single play element. Similarly, *Retreating Backwards* and *Retreating* were closely connected. A lot of similarity between elements can be explained by their frequent co-occurrence – for example, *Retreat* and *Bipedal* showed high similarity because chimpanzees often retreat from the play partner while bipedal.

**Figure 3:**
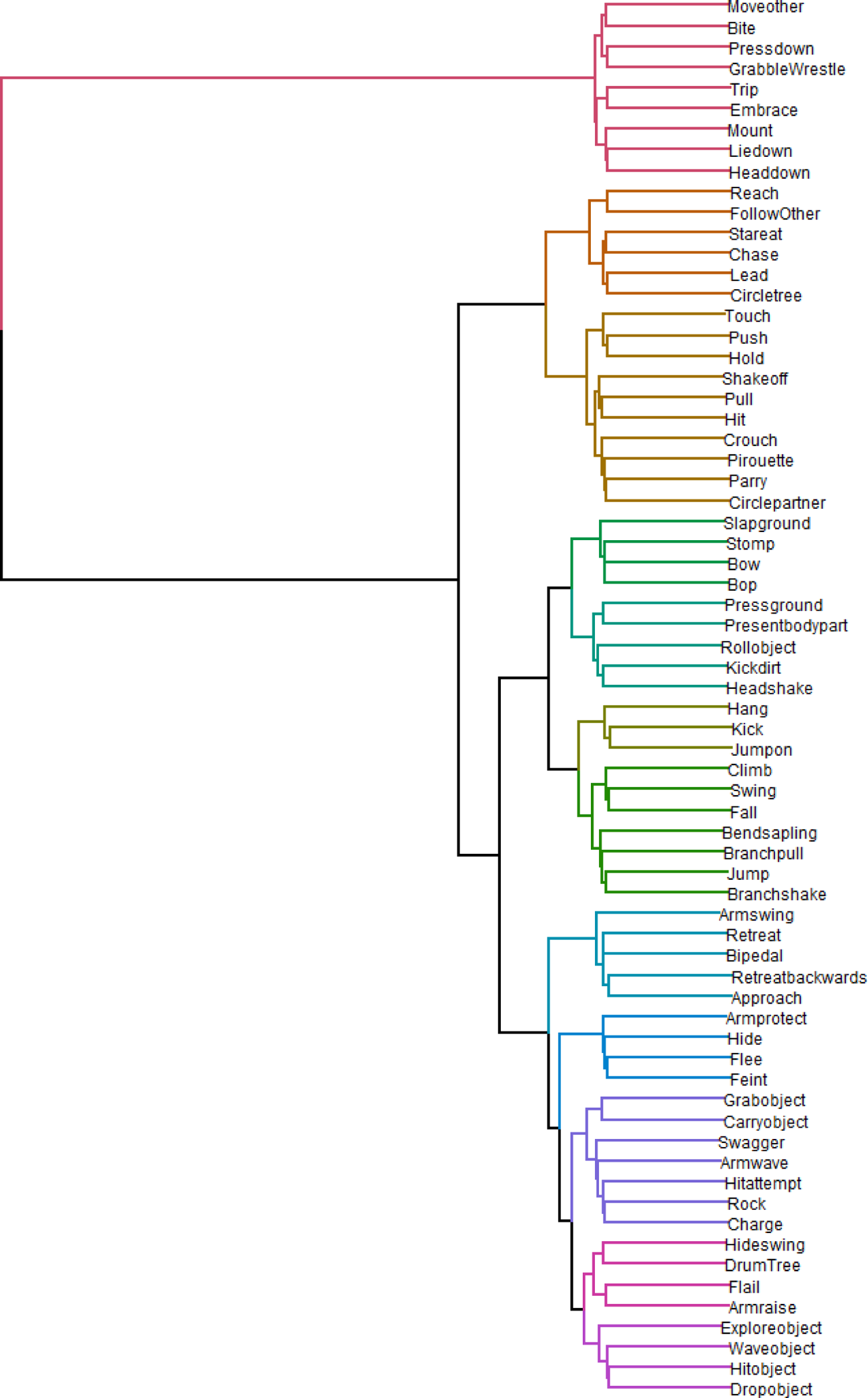
Dendrogram of hierarchical clustering of distances between play elements. Branch colours indicate established cluster membership. Optimal cluster solution: 11 clusters.

**Table 2:**
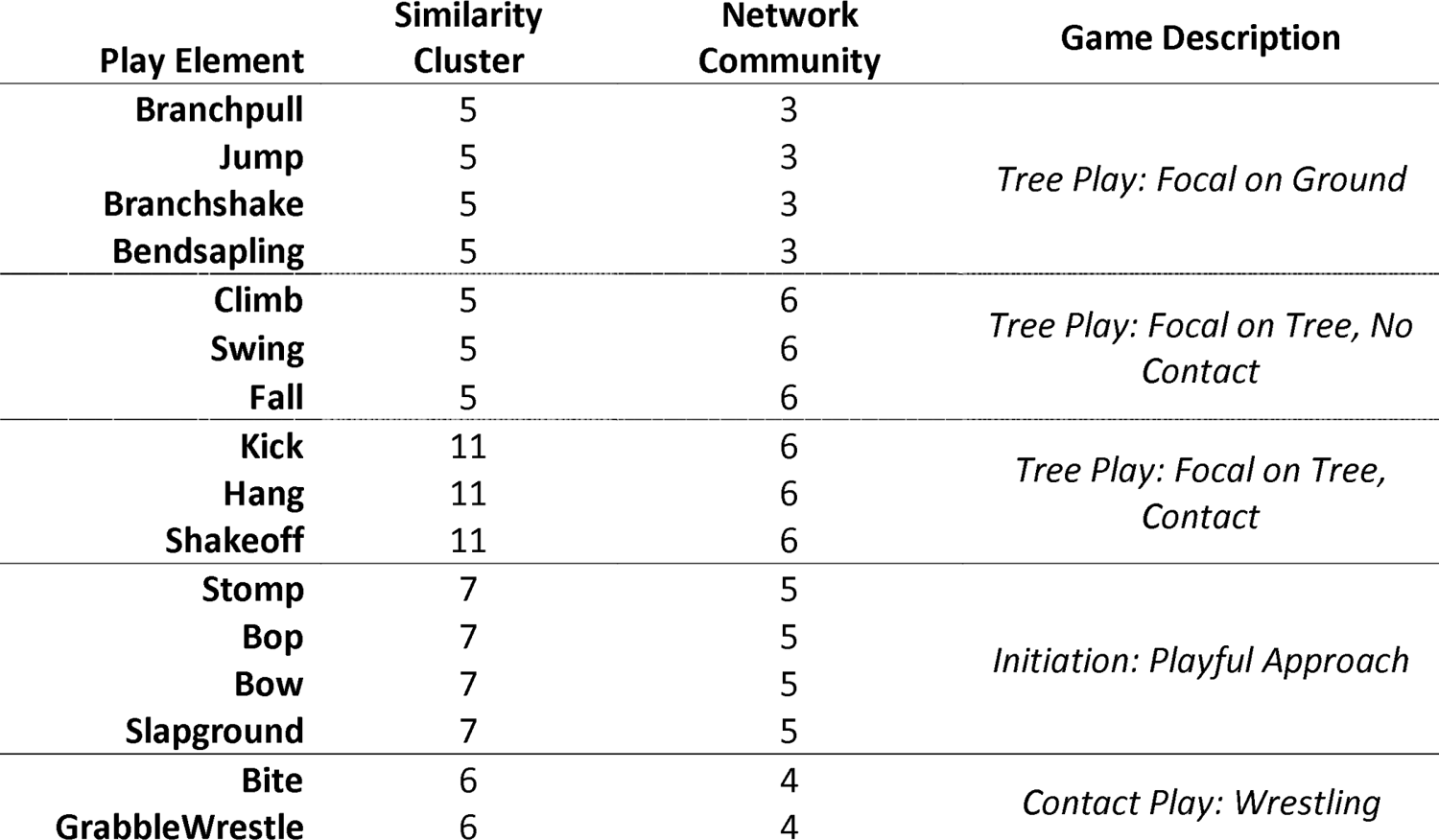
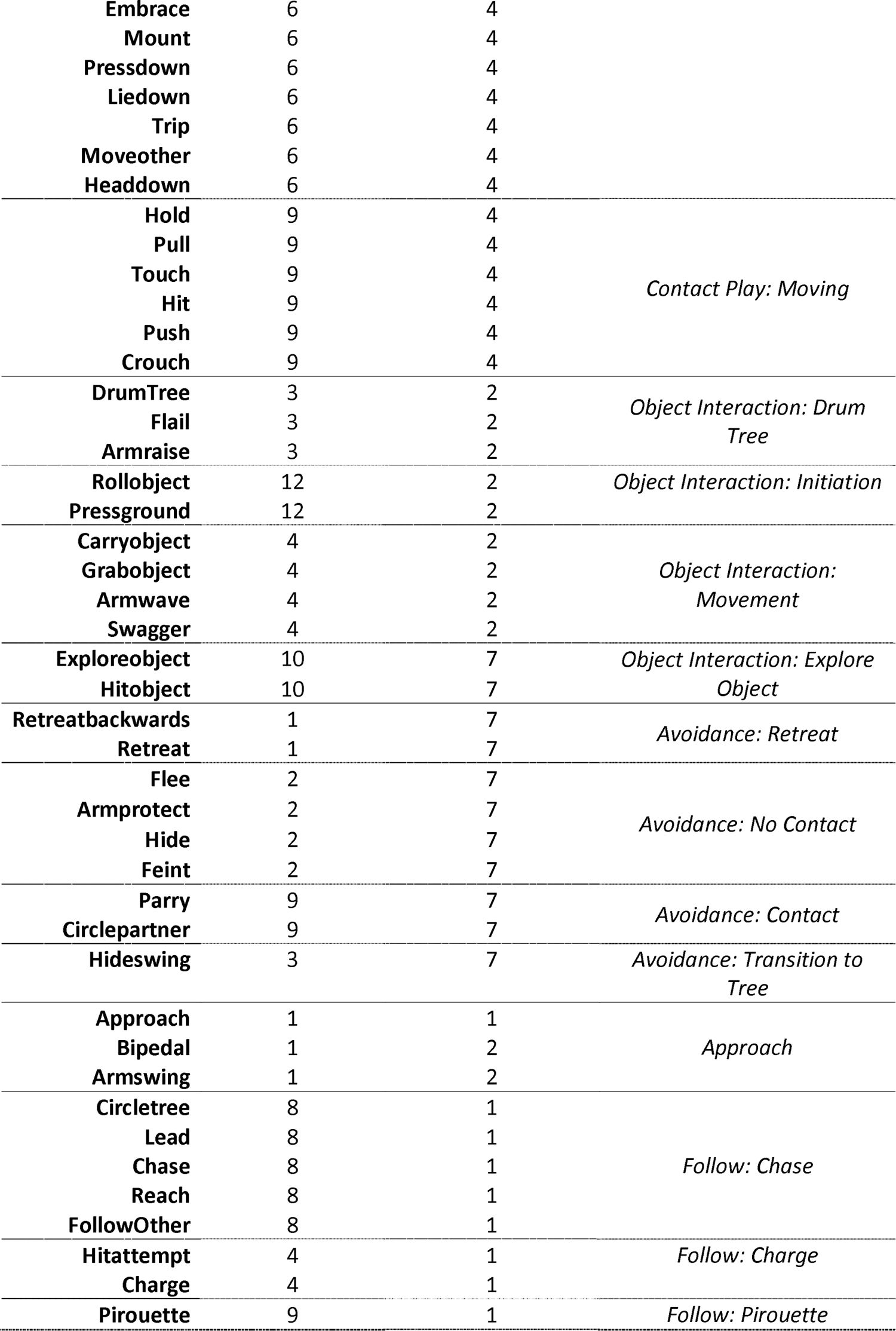
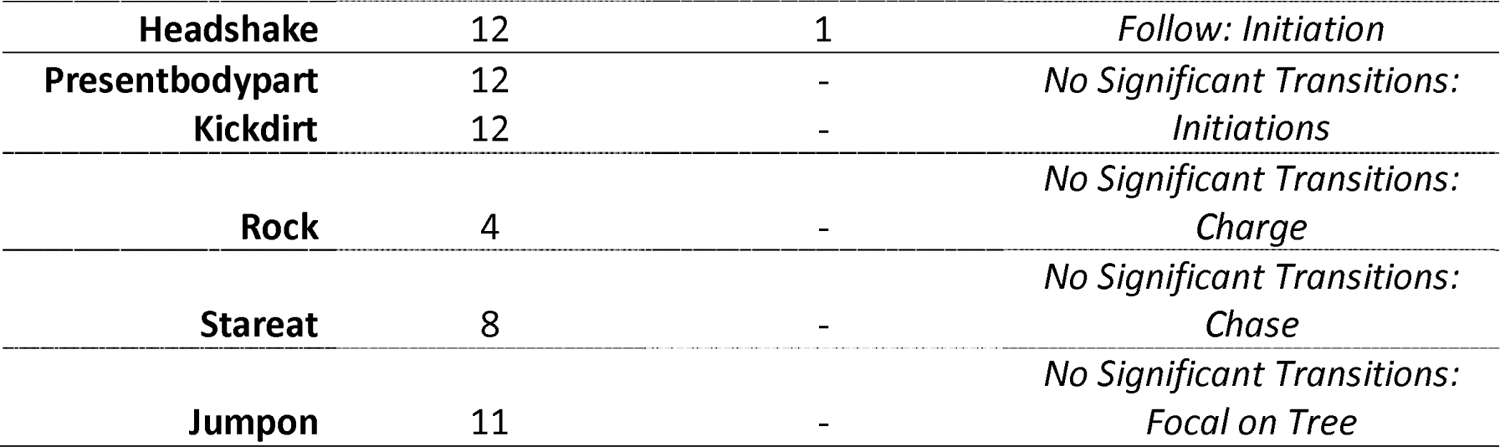
Play elements with their cluster/community assignment for both the similarity and network of transition probabilities

### d) Network structure

In contrast to the similarity clusters, which assess whether two elements are used at similar points in a sequence, the transition network (Figure 4) describes which consequent follows which antecedent. The network only depicts transitions that occurred at higher-than-expected rates and occurred at least 5 times in the dataset. Colours indicate community membership. As the high modularity of the network community detection algorithm (modularity = 0. 65) indicates, there were seven clearly distinguished communities in the network. If community assignment was random, we would expect around 32% of transitions between the elements within communities, but we observed 48% of transition within communities – a 1.5-fold increase. Connections between communities were often due to elements that can be used in different situations. For example, *Shake Off* is used when playing *wrestling* with a partner to get away, but equally when the player is *hanging* off a branch or *retreating* – hence, the element is connected to three communities. Individuals *stomp* when initiating play in combination with *Bop* and *Bow*, but also when they were *bending* a small tree and holding onto it.

**Figure 4:**
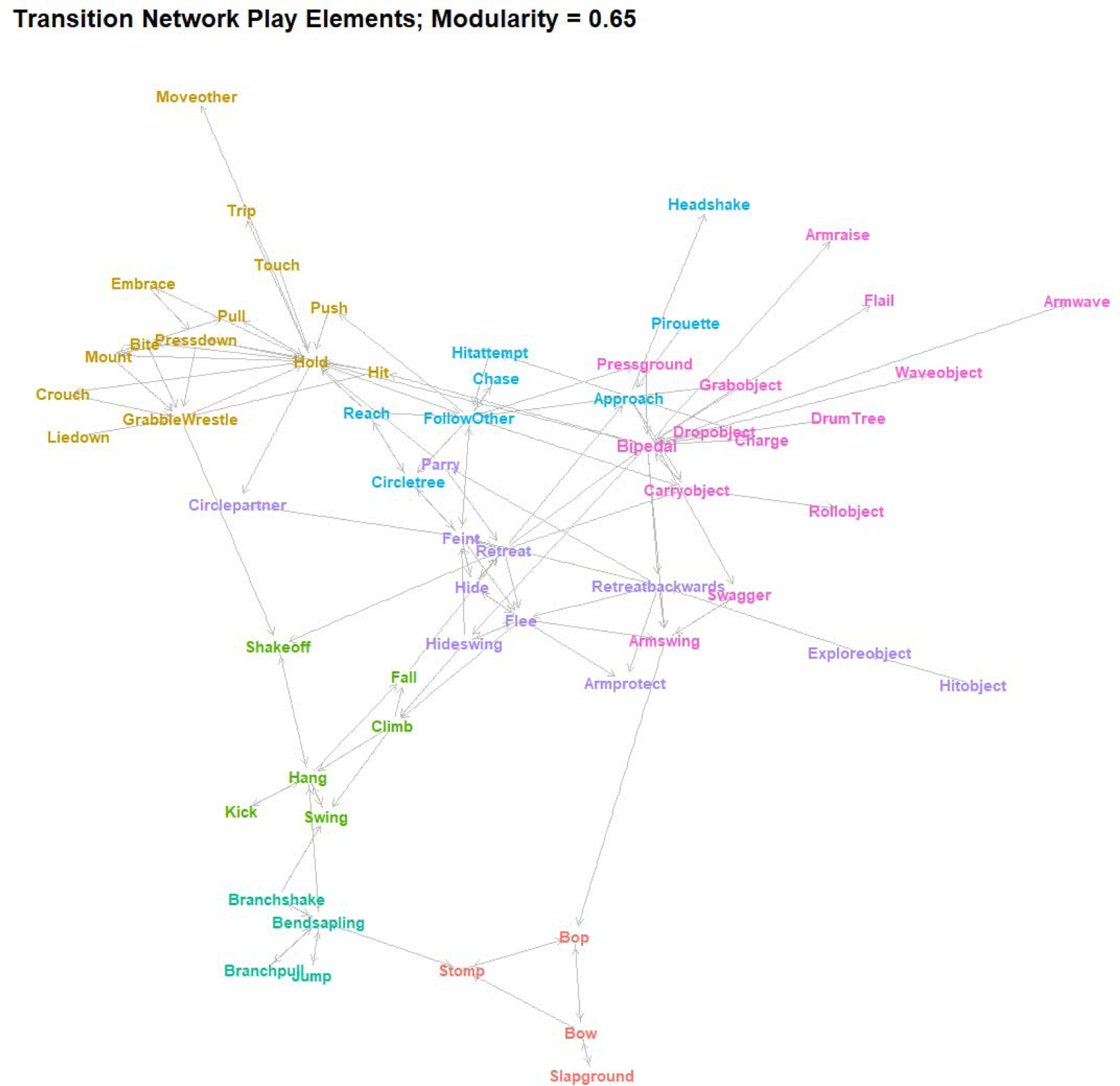
Network plot of weighted transition probabilities between play elements. Play elements are nodes, significant transitions that occur at least 5 times are edges (directed), and colour indicates cluster membership.

For the interpretation of communities, in combination with the similarity clusters, see Table 2. There was considerable overlap between the two approaches, with small variation arising mainly because several elements did not have any significant transitions above threshold level, and the combination of object-related and movement elements resulted in overlap between the chase and object clusters. The different cluster combinations (‘games’) can be categorised broadly by whether they involved climbing by either partner, had physical contact, involved chasing, involved objects, or were play invitations. For the latter, one clear cluster emerged, consisting of *Bop*, *Bow*, *Stomp*, and *Slap Ground*, which individuals often combined and repeated in quick succession to indicate that they were willing to play. Some other, rarer elements (*Present Body Part*, *Rock*, *Kick Dirt*, *Stare At*) can fulfil a similar function. Play elements routinely used when one or both individuals were in a tree transitioned into each other at high rates, depending on the role of the focal individual. When the player was on the ground and the partner in the tree, individuals would often *bend the tree* (the most central element of this cluster), and then *pull* or *shake* it, sometimes while *jumping*. While players were in the tree, they *climb* up and then *hang* while *swinging*, *kicking* the partner, *shaking them off*, and ultimately *falling*.

Most contact play formed one large community in the network, with elements transitioning into each other at high rates. Based on the similarity of transition probabilities, we could differentiate two groupings: contact play that involves players to stay in one spot (*Bite*, *Wrestle*, etc), centred on *holding* the partner in place; and those that involve one player trying to get away from their partner while still in contact (*Push, Trip* etc).

The different object-related play elements were connected, including detached objects and trees. Chimpanzee players held onto objects once they had grabbed them and then manipulated them in different ways. Object contact was the defining element of this type of play. A common way for the Bossou chimpanzees to initiate play with object contact was to *roll objects* towards the partner or *press the ground*. Individuals will often *drum trees*. After, players regularly *hide swing* (swinging around a tree at speed), which will lead to tree play after. Players *wave objects* about while *swaggering* towards the partner and *flailing* or *waving* their arms. Many of the social object elements were connected to *retreating* movements, with the player retreating while holding an object, which explains the community overlap of object interactions and avoidance movements.

The remaining cluster combinations were related to chasing play on the ground. Again, we can identify different roles of the player: One community were those elements strongly connected to movements used to avoid the partner *retreating* or *retreating backwards* from them and *hiding* behind trees or *feinting* directional changes, often lifting their *arm protectively*. They will *circle the partner* while *parrying hits*. The last cluster combination involved the opposite, with the individual *approaching* the partner (sometimes following a *pirouette* as play initiation, often combined with *bipedal* movements and *arm swings*), *chasing*, and trying to make physical contact while the partner flees (*Reach*, *Hit Attempt*).

## Discussion

In this study, we explored the sequence structure of Western chimpanzee play behaviour for the Bossou community. We were interested in how predictable play was, and whether we find distinct ‘games’ with clear rules for sequences used by each player. Despite the large number of play elements and of transitions that were observed infrequently, only a small number of transitions occurred at higher-than-expected rates. Information about the preceding play element allowed for more accurate predictions than random choice, and the predictions became more accurate when including more antecedent elements – however, play retained its unpredictability, as the accuracy of predictions remained low. The reason for this can be found in the patterns of different ‘games’: we showed that there were several clusters of highly connected play elements with similar transition patterns. Thus, when a player was climbing in a tree, there were only few play elements available to them, but the exact order cannot be predicted. This appears to be similar to human games – if two children play tag, there is a finite number of play elements that each of them can use to keep the game going, but it is not in either players interest to let the partner know which one is next. Importantly, the clusters we detected indicated clear roles for at least some of the games, with play partners on the ground acting different from the one in the tree and avoiding play elements clearly distinguished from approaching elements in chases.

Animal play behaviour is characterised by its unpredictable nature compared to other contexts, leading to theories that it has evolved as a method for young individuals to learn social and motor skills that will become important later in life (Fagen, 1981; Smith, 1982). We show that, at least for chimpanzees, play is a mix of predictability and unpredictability – while knowledge of previous actions allows us to improve predictive accuracy, play sequences are not simple Markov chains, where one or few antecedent actions allow for accurate reactions. However, that does not mean that play is random, as clear games emerged from our bottom-up, data driven approach. We detect clusters of elements that are used together and interchangeably, indicating a rule-based system were the game limits the number of appropriate responses. Further studies will have to determine whether non-linear prediction methods, e.g., deep learning (Chollet & Allaire, 2018) could increase predictive accuracy, and whether action sequences are better described using non-Markov processes (Kershenbaum et al., 2014). Using a naïve Bayes classifier strongly improved predictive accuracy, and more complex machine learning algorithms and a larger dataset could further extend our ability to detect transition patterns. For now, this study demonstrates the power of a ‘grammar of actions’ approach (Pastra & Aloimonos, 2012), where methods from natural language processing and syntactical analysis are employed to understand the sequential nature of behavioural actions in humans and non-human animals. Our study presents evidence that the ability of chimpanzees to produce hierarchically structured sequences is not limited to their communication (Arcadi, 1996; Girard-Buttoz et al., 2022; Liebal et al., 2004) and tool-related behaviour (Carvalho et al., 2008; Estienne et al., 2017; Vale et al., 2017), but is also prevalent in fast-paced social interactions that require adaptation to multiple partners in real time (McCarthy et al., 2013).

Some of the games have previously been identified by researcher when coding primate play – for example, many studies code ‘rough-and-tumble’ play as an overarching category for all physical play in close contact (Palagi et al., 2016). Our results show that this category can be established with a data driven approach. The same is true for chasing games. Another overarching context is tree-related play, either with the player climbing or on the ground. Lastly, we identified social object play as its own context, which equally has attracted research in the past as a possible window into game-like behaviour (Shimada, 2006; Tanner & Byrne, 2010). Each of those games consisted of some central elements – holding the partner, moving towards them, moving away from them, holding an object, hanging from a tree etc. – that defined the context, with other elements added more freely, similar to tool use sequences in chimpanzees (Carvalho et al., 2008). We found clear evidence of role-reversal between players, as has long been described for play across species (Fagen, 1981) – players on the ground have a clear role in tree play that differs from those of the partner in the tree, and chasing players use different play elements than those fleeing. However, it needs to be kept in mind that the small sample size for many of the elements makes some of these clusters unreliable and dependent on researcher choices for the UMAP and clustering algorithms.

The specific research context of this study, using video footage of the Bossou chimpanzees while they are in the forest clearing of the outdoor lab, constrains the number of different games that could be observed – for example, water games as in mountain gorillas (Costa et al., 2019) cannot be observed in this environment. The physical substrate around the outdoor lab limits the amount of arboreal play. Thus, while we describe a method to detect games, larger datasets and more varied collection contexts would be necessary to characterise chimpanzee games more broadly. We are not trying to describe species-specific play patterns for chimpanzees in general (which probably include strong developmental, individual, dyadic, and group-level effects), but show that in this fairly standardised sample, chimpanzee play shows complex sequential patterns. Importantly, our approach would allow direct comparisons between different communities of chimpanzees, based on transition probabilities and network patterns. As the form of play is less defined by its function than for example tool use, this might be a useful approach to study cultural differences in a meaningful way (Boesch et al., 2020).

One aspect currently missing from the picture is partner behaviour: while within-player behaviour shows limited predictability, it might be more predictable when knowing what the partner did. Chimpanzees and other primates engage in turn-taking when communicating (Chow et al., 2015; Fröhlich, 2017), and play has been described as a context that elicits joint commitment between players, with clear evidence that they re-establish that commitment after breaks (Heesen, Bangerter, et al., 2021; Heesen, Zuberbühler, et al., 2021). Thus, we need an approach that understands social interactions (including play) as a complex system of decisions taken by all involved individuals. One question is whether play is indeed more complex in its sequential structure than other social contexts, such as grooming or aggressions, or communicative exchanges. The statistical analyses underlying this study can be replicated using any data consisting of sequences of discrete elements.

The data collection, pre-processing, and analytical choices of this study introduce several researcher degrees of freedom that limit generalisability of results (Wicherts et al., 2016). Thus, we are interpreting all results regarding the structure of play element transitions conditional on the coding scheme and group. The Bossou chimpanzees are a very small group and subadult players lack same-aged play partners. Results were based on a small number of players who had an outsized impact on the dataset, and accounting for individual-level idiosyncrasies and age-dependent contingencies in transitions between play elements might dramatically improve predictability (Cordoni & Palagi, 2011). Many play elements were rare, and we had to make choices on how to combine them; there was still considerably uncertainty for some of the transition probability estimates. We set strict cut-offs for significance levels and the minimum number of observed transitions to err conservatively, but an increased dataset or different thresholds might influence results. Another choice we had to make was regarding co-occurring play elements. We chose to use permutations to randomly assign which elements occurred at what point in the sequence, but this approach necessarily increases noise in the data. Lastly, every study of play behaviour is using a different ethogram, with different levels of complexity. We would predict that a simpler coding scheme would result in higher predictability. Because of the complexity of the coding scheme chosen here, no inter-rater reliability was performed, thus results should be interpreted as conditional on the coding process.

In summary, we show that chimpanzee play behaviour is a complex sequential process with an identifiable hierarchical structure – chimpanzees play games consisting of play elements that are interchangeable in their sequence position and transition into each other at higher rates than they transition into play elements that are representative of other games. Information about previous actions allows for prediction of subsequent elements and including more antecedent elements improves accuracy. Our results show that there is considerable leeway to study decision-making and cognitive complexity in animal social interactions on the micro-level (Gygax et al., 2021), but this process, like the study of communication, requires detailed video analysis of long-term data (Hobaiter & Byrne, 2011). In the future, being able to achieve reliable behaviour recognition from video databases, as has been demonstrated for the Bossou chimpanzees (Bain et al., 2021), could be a valuable tool in reducing the coding effort involved. As it stands, our results further highlight the special place play behaviour holds in the cognitive and behavioural development of chimpanzees – by creating a save environment to explore and train fast-paced behavioural sequences, it allows young individuals to learn to predict how a partner will react in different social situations.

## Data Availability

All data and R scripts are available in a bespoke GitHub repository that allows reproduction and replication (https://github.com/AlexMielke1988/Mielke-Carvalho_Chimpanzee-Play).

## Acknowledgements

This research was funded by the British Academy through AM’s Newton International Fellowship. AM also received funding from the Leverhulme Trust. We thank Professor Tetsuro Matsuzawa for the long-term efforts to develop and support the research on wild chimpanzees, at the Bossou Field Station, in Guinea. Those efforts were supported by grants from MEXT (#12002009, #16002001, #20002001, #24000001, #16H06283) and JSPS (Core-to-core CCSN and U04-PWS). We thank Daniel Schofield and Misato Hayashi for their continuous help in curating the Bossou video archive, and Sophie Berdugo for help identifying the chimpanzees. We are also very grateful to Dora Biro, Catherine Hobaiter, David R. Braun, and all the KUPRI researchers who helped to collect field data at Bossou between 2009 and 2013. Special thanks are due to Direction General de la Recherche Scientifique et l’innovation Technologique (DGERSIT) and to Dr. Ali Gaspard Soumah and the Institut de Recherche Environnementale de Bossou (IREB), République de Guinée, for assistance and research permission to conduct field work at Bossou; as well as research assistants Boniface Zogbila, Gouanou Zogbila, Henry Didier Camara, and Gilles Doré, Pascal Goumy, for their invaluable assistance in the field.

